# Snf2 controls pulcherriminic acid biosynthesis and connects pigmentation and antifungal activity of the yeast *Metschnikowia pulcherrima*

**DOI:** 10.1101/494922

**Authors:** Deborah Gore-Lloyd, Inés Sumann, Alexander O. Brachmann, Kerstin Schneeberger, Raúl A. Ortiz-Merino, Mauro Moreno-Beltrán, Michael Schläfli, Pascal Kirner, Amanda Santos Kron, Kenneth H. Wolfe, Jörn Piel, Christian H. Ahrens, Daniel Henk, Florian M. Freimoser

## Abstract

*Metschnikowia pulcherrima* synthesizes the red pigment pulcherrimin, from cyclodileucine (cyclo(Leu-Leu)) as a precursor, and exhibits strong antifungal activity against notorious plant pathogenic fungi such as *Botrytis* and *Gibberella* (i.e., *Fusarium*). This yeast therefore has great potential for biocontrol applications against fungal diseases; particularly in the phyllosphere where this species is frequently found. To elucidate the molecular basis of the antifungal activity of *M. pulcherrima*, we compared a wildtype strain with a spontaneously occurring, pigmentless, weakly antagonistic mutant derivative. Whole genome sequencing of the wildtype and mutant strains identified a point mutation that creates a premature stop codon in the transcriptional regulator *SNF2* in the mutant strain. Complementation of the *snf2* mutant strain with the wildtype *SNF2* gene restored pigmentation and recovered the strong antifungal activity of *M. pulcherrima* against plant pathogens *in vitro* and on cherries. Ultra-performance liquid chromatography-high resolution heated electrospray ionization mass spectrometry (UPLC HR HESI-MS) proved the presence and structure of the pulcherrimin precursors cyclo(Leu-Leu) and pulcherriminic acid and also identified new compounds that likely represented an additional precursor and degradation products of pulcherriminic acid and/or pulcherrimin. All of these compounds were identified in the wildtype and complemented strain, but were undetectable in the pigmentless *snf2* mutant strain. These results thus identify *SNF2* as a regulator of antifungal activity and pulcherriminic acid biosynthesis in *M. pulcherrima* and provide a starting point for deciphering the molecular functions underlying the antagonistic activity of this yeast.

**Significance statement:** *Metschnikowia pulcherrima* is a strongly antifungal yeast and a most promising species for the control of notorious plant diseases. This multidisciplinary study on the *M. pulcherrima* mode of action compared a wildtype isolate with a pigmentless mutant exhibiting reduced antifungal activity. The transcriptional regulator Snf2 was identified as a “biocontrol regulator” controlling antifungal activity of *M. pulcherrima* via *PUL* gene transcription, cyclodipeptide synthesis and additional, yet uncharacterized mechanisms. The identification of cyclo(Leu-Leu), pulcherriminic acid, as well as novel precursor and degradation products of pulcherrimin, opens up new avenues for research on the metabolism and functions of pulcherrimin. Overall, this works establishes *M. pulcherrima* as a genetically tractable model and will benefit the development of biocontrol solutions for important plant diseases.

## Introduction

The yeast *Metschnikowia pulcherrima* is globally distributed and frequently isolated from the phyllosphere; in particular from flowers and fruits (1-3). Competition assays of 40 yeasts against a diverse set of 16 filamentous fungi identified *M. pulcherrima* as the overall strongest antagonist (4) and the same isolate was highly competitive against other antifungal yeasts on apples (5). The species exhibits strong antagonistic activity against apple postharvest diseases caused by *Alternaria, Aspergillus, Botrytis, Fusarium, Monilinia*, and *Penicillium* species (6-11) and was also identified as a potential biocontrol organism against food-borne pathogens of freshly cut apples (12) as well as fungal grape diseases (13, 14). Besides fungal pathogens of fruits, *M. pulcherrima* also inhibits other yeasts (15, 16). In addition to biocontrol applications, *M. pulcherrima* is being considered for the synthesis of the fragrance molecule 2-phenylethanol (17), for biodegradation of the mycotoxin patulin (18), and for biofuel production (19), and is being evaluated for the production of wines with lower alcohol content (20, 21).

The designation “the most beautiful *Metschnikowia*” (translation of the Latin word “*pulcherrima*”) is based on the cells’ uniform and spherical appearance in microscopic preparations (22), and not the red pigment produced under certain growth conditions. However, the red pigment, forming spontaneously by a non-enzymatic reaction in the presence of iron under certain culture conditions, is a remarkable characteristic of *M. pulcherrima* colonies (23). The compound responsible for this phenotype is described as pulcherrimin; an iron (III) chelate of pulcherriminic acid (2,5-diisobutyl-3,6-dihydroxy-pyrazine-1,4-dioxide). Pulcherriminic acid is formed by the oxidation of cyclodileucine (cyclo(Leu-Leu) (24, 25). Pulcherrimin was first described in yeasts more than 60 years ago, but only very recently the four genes involved in its biosynthesis and transport in *Kluyveromyces lactis* have been identified (26), and the structure of the molecule synthesized by yeasts has not been determined by modern analytical methods. However, pulcherrimin is also produced by certain bacteria such as *Bacillus licheniformis*, where the two genes *yvmC* and *cypX*, coding for a cyclodipeptide synthase and cytochrome P450 oxidase, respectively, are responsible for its biosynthesis (27-29). Expression of *yvmC* and *cypX* and the formation of pulcherrimin are intricately regulated by factors such as growth stage, iron availability, or environmental stress (30, 31). Iron chelation by pulcherriminic acid and formation of insoluble pulcherrimin, causing iron depletion in the growth substrate, is thought to mediate antimicrobial activity of bacteria and yeasts (9, 16, 30, 32, 33). However, alternative hypotheses to this mode of action cannot be excluded (e.g., antimicrobial activity due to precursors and independent of iron, regulatory effect on plant, fungal or bacterial responses, defense reaction against toxic iron levels) (22, 33).

In the present study, we characterized the mode of action of *M. pulcherrima* by comparing pigmentless mutants with reduced antifungal activity with their wildtype progenitor. Comparison of the mutants with the *de novo* assembled genome of the wildtype strain allowed identifying a premature stop codon in the transcriptional regulator *SNF2* as the cause for the observed phenotypes, which was confirmed by developing genetic tools and complementation of the mutant strain. We show that the *snf2* mutant neither synthesized pulcherriminic acid or cyclo(Leu-Leu), two precursors of pulcherrimin, nor any other precursor or degradation product of pulcherriminic acid. This phenotype was likely caused by strongly reduced transcript levels of the pulcherrimin biosynthesis genes *PUL1* and *PUL2 (26)*. These results establish the role of *M. pulcherrima SNF2* in the regulation of antagonistic activity and pulcherrimin biosynthesis. Our findings on pulcherrimin metabolism are not only biotechnologically relevant, but will allow further defining the mode of action of *M. pulcherrima*. Furthermore, the tools established in the course of this work open up new opportunities for application-oriented, fundamental research on the *M. pulcherrima* antifungal activity that will benefit phytopathology and the development of biocontrol solutions for important plant diseases.

## Results

### Naturally occurring, pigmentless *M. pulcherrima* mutants are less antagonistic than the wildtype

Based on the original literature, *Metschnikowia pulcherrima* synthesises pulcherriminic acid from cyclodileucine (24). In the presence of iron (III), a non-enzymatic reaction converts pulcherriminic acid into the red pigment pulcherrimin. Pigmentless, white mutants have previously been reported after nitrosoguanidine treatment (33) and were here found to occur spontaneously among freeze-dried *M. pulcherrima* cells that were stored at 22 °C for several months. Since pulcherrimin has been implicated in the antagonistic activity of *M. pulcherrima* and *Bacillus licheniformis*, supposedly due to its iron immobilizing activity (30, 33), we characterised pigmentless *M. pulcherrima* mutants with respect to their antagonistic activity against plant pathogenic fungi.

The *M. pulcherrima* wildtype isolate APC 1.2 (WT) exhibited increased pigmentation with increasing iron concentrations, while the mutant cells (isolate W8) remained pigmentless (white) irrespective of the iron concentration in the growth medium (Figure 1A). Binary competition assays on potato dextrose agar (PDA) plates (Figure 1B) confirmed strong inhibition of the plant pathogenic fungus *Botrytis caroliniana* by the *M. pulcherrima* WT (98% reduction in the growth area as compared to a control growing in the absence of the yeast) (Figure 1C). In contrast, the *M. pulcherrima* pigmentless mutant W8 inhibited *B. caroliniana* less than the wildtype (80% reduction of the growth area). This reduced antifungal activity of the non-pigmented mutants was similarly observed in competition assays against *Gibberella fujikuroi* and *Fusarium oxysporum* (56% and 45% vs. 89% and 81% reduction) and with two other, independent, pigmentless *M. pulcherrima* mutants W10 and W11 (37%-51% reduction vs. 75% reduction by the wildtype) (Supplementary Figure S1). The strong antagonistic activity of the *M. pulcherrima* WT and the reduced antifungal effect of the W8 mutant were confirmed in bioassays on cherries that were infected with *B. caroliniana* (Figure 1D). Compared to cherries infected with the plant pathogen alone, the *M. pulcherrima* WT strongly suppressed *Botrytis* on cherries with respect to rot diameter and abolished mycelium development (Figure 1E). In contrast, the W8 mutant only weakly suppressed *Botrytis*, leading to strongly increased rot diameter and mycelium development of *B. caroliniana,* albeit in lower frequency and less profusely than in the cherries only inoculated with *Botrytis* (Figures 1D, E). Overall, these results demonstrated reduced (but still detectable) antifungal activity of the pigmentless, white *M. pulcherrima* mutant W8 *in vitro* as well as in cherries.

**Figure 1.**
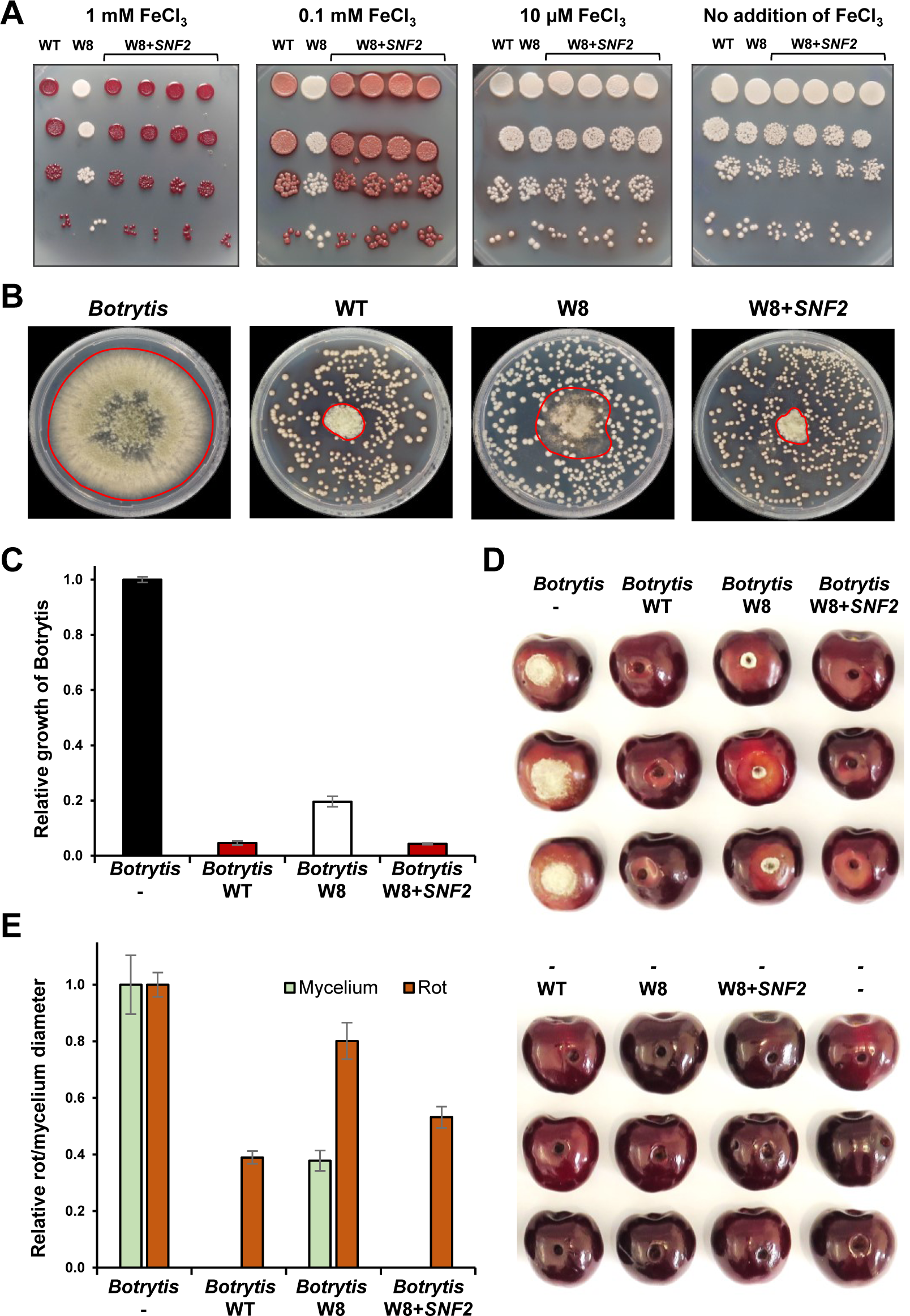
Complementation with *SNF2* restores pulcherrimin biosynthesis and antagonistic activity in a pigmentless *M. pulcherrima* mutant. All experiments were performed with the *M. pulcherrima* wildtype (WT; isolate APC 1.2), mutant (W8) and complemented (W8+*SNF2*) strains. A) The pigmentless *M. pulcherrima* mutant (W8, second column) remained white irrespective of the amount of iron (III) (FeCl_3_; 0-1 mM) added to the growth medium, while the wildtype (WT, first column) and four complemented strains (columns 3-6) showed strong pigmentation, indicative of pulcherrimin biosynthesis, if grown in medium supplemented with at least 100 μM FeCl_3_. B, C) Binary competition assays against *Botrytis* (*B. caroliana*; isolate EC 1.05, SH177344.07FU) were performed on PDA plates and the relative growth area, as compared to the wildtype, was quantified. The area occupied by *Botrytis* mycelium is marked in the photograph and the data in the histogram show the means and standard errors of four replicates. D, E) Bioassays with freshly harvested, conventionally grown cherries were performed with *Botrytis* as the plant pathogen. The diameter of the *Botrytis* mycelium developing around the artificially introduced lesion, as well as the (larger) diameter of the rot (lighter colored, sunken area around the lesion) were measured. The *M. pulcherrima* wildtype and complemented strains abolished the formation of *Botrytis* mycelium around the lesion. None of the yeasts alone, without *Botrytis*, caused rot or mycelium development on the cherries.

### Pigmentless *M. pulcherrima* mutants do not secrete cyclodileucine or pulcherriminic acid

The white *M. pulcherrima* mutant seemed to lack the red pigment pulcherrimin, but it was not clear if the pigment produced by our *M. pulcherrima* isolate APC 1.2 indeed corresponded to the compound described as pulcherrimin, if the precursors cyclo(Leu-Leu) and/or pulcherriminic acid were not synthesized, or if another defect prevented the formation of the iron chelate. Therefore, we analysed the culture supernatants of wildtype and mutant *M. pulcherrima* cells by ultra-performance liquid chromatography-high resolution heated electrospray ionization mass spectrometry (UPLC HR HESI-MS) in order to identify the soluble pulcherrimin precursors cyclo(Leu-Leu) and pulcherriminic acid.

High resolution MS measurements in combination with ^15^N- and ^13^C-labelling experiments allowed determining the molecular formula. Metabolites secreted by the *M. pulcherrima* WT and W8 mutant during growth were absorbed with Amberlite^®^ XAD16 resin and extracts thereof were analysed by UPLC HR HESI-MS. We identified at least five dipeptides that were secreted by *M. pulcherrima* WT cells, but were not detected in the W8 mutant supernatants (Figures 2A, B). Comparison with a cyclo(Leu-Leu) standard identified **1** (227.176 *m*/*z* [M+H]^+^, C_12_H_22_N_2_O_2_) as cyclo(L-leucyl-L-leucyl) (Supplementary Figure S2). Extensive UPLC HR HESI-MS analyses, including reverse stable isotope labeling with L-leucine, L-isoleucine and L-valine in a ^13^C-background, confirmed the incorporation of two L-leucine into all five compounds (Supplementary Figure S3). Based on these analyses, **2** (257.150 *m*/*z* [M+H]^+^, C_12_H_20_N_2_O_4_) was identified as pulcherriminic acid and **3** (259.165 *m*/*z* [M+H]^+^, C_12_H_22_N_2_O_4_) might represent a di-*N*-hydroxylated intermediate of **1** (Figure 2A, Supplementary Figures S3, S4). The molecular formula of **4** (275.161 *m*/*z* [M+H]^+^, C_11_H_20_N_2_O_2_) suggests an additional hydroxylation of **2** and also ring opening, whereas **5** (229.155 m/z [M+H]^+^, C_11_H_20_N_2_O_2_) indicates a decarboxylation event due to the incorporation of only one C_6_- and one C_5_-L-leucine derived carbon backbone, instead of two C_6_-carbon skeletons. In this respect, **5** might represent a degradation product of **4** (Figure 2A, Supplementary Figures S3, S4). Besides that, we observed two C_4_-backbones in all compounds after L-valine supplementation. Here, valine is likely metabolized via 2-isopropylmalate into leucine before incorporation (Figure 2A, Supplementary Figures S3, S4). Based on our results, we identified three of these compounds as the pulcherrimin precursors cyclo(L-leucyl-L-leucyl), pulcherriminic acid, and a yet uncharacterized precursor or shunt product, while the other two compounds likely represent novel degradation products.

**Figure 2:**
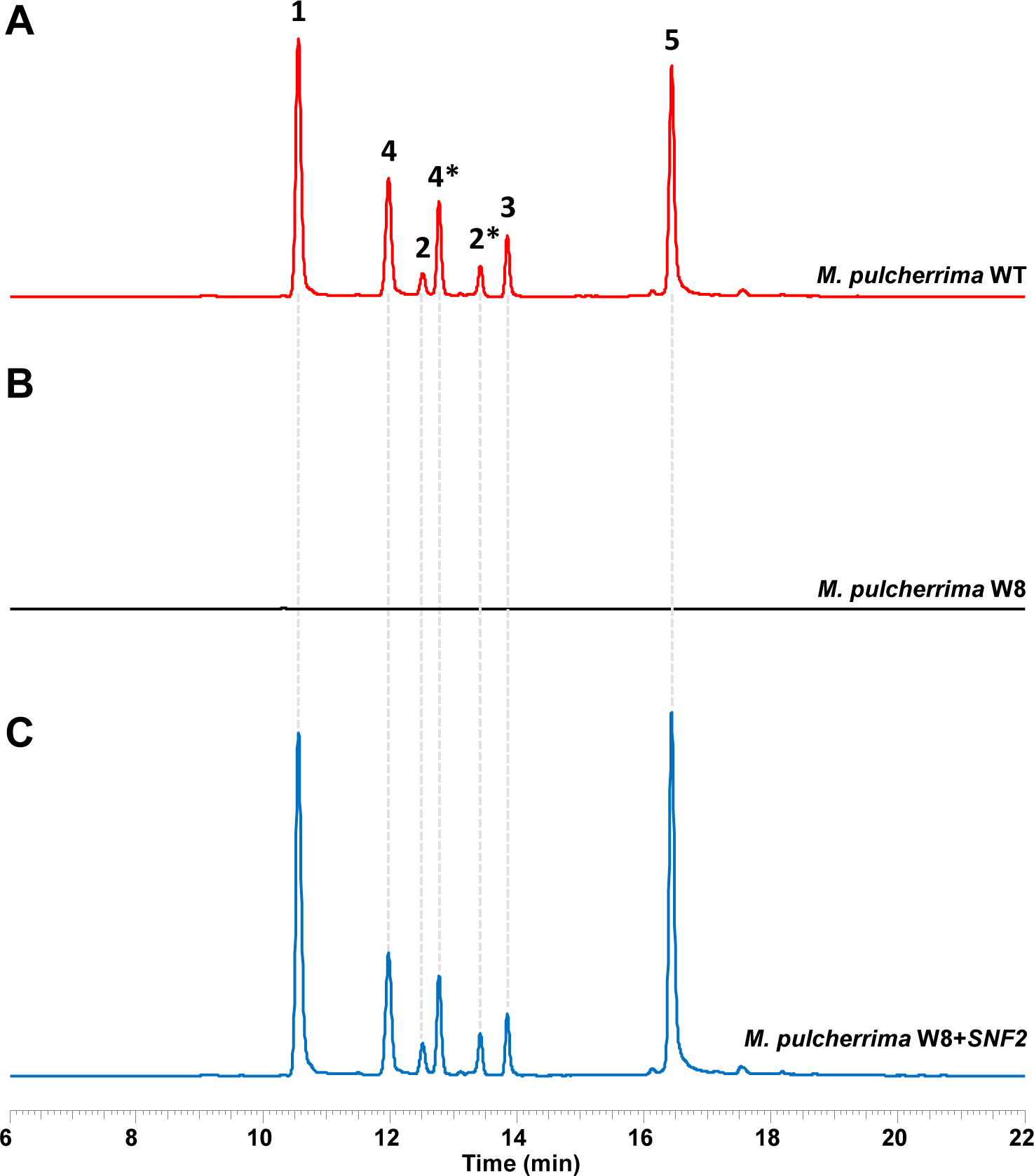
A pigmentless *M. pulcherrima* mutant does not secrete cyclic dipeptides. UPLC HR HESI-MS metabolic profiles for (1, cyclo(Leu-Leu, 227.176 *m*/*z* [M+H]^+^, C_12_H_22_N_2_O_2_)) derived derivatives in the wildtype of *M. pulcherrima* (top, red), *snf2* mutant (middle, black) and a *SNF2* complemented mutant (bottom, blue). (2, pulcherriminic acid, 257.150 *m*/*z* [M+H]^+^, C_12_H_20_N_2_O_4_), (3, 259.165 *m*/*z* [M+H]^+^, C_12_H_22_N_2_O_4_), (4, 275.161 *m*/*z* [M+H]^+^, C_12_H_22_N_2_O_5_), (5, 229.155 *m*/*z* [M+H]^+^, C_11_H_20_N_2_O_2_)). Potential tautomers are marked with an asterisk.

### Pigmentless *M. pulcherrima* mutants harbor a point mutation leading to a premature stop codon in the gene encoding the transcriptional regulator Snf2

In order to identify the underlying genetic mutations leading to the lack of cyclodipeptide biosynthesis and weakly antagonistic *M. pulcherrima* phenotype, a high-quality reference genome for the *M. pulcherrima* wildtype strain APC 1.2 was *de novo* assembled relying on long-read PacBio sequencing.

The final, assembled and annotated genome (assembly with Canu (34), 254x average coverage) indicated a genome size of 15.88 Mbp, a G+C content of 46%, and 5922 protein-coding genes (Supplementary Table S1). The genome contains two genes for tRNA-Ser(CAG), consistent with a CUG-Ser genetic code as expected for this species, which is relatively closely related to *Candida albicans*. The nuclear genome was assembled into 7 large scaffolds (>600 kb). Six of the 14 ends of these 7 scaffolds terminated at 18S or 26S ribosomal RNA genes, and the others terminated at putative subtelomeric repeats that were shared among scaffold ends. The 5S rRNA genes are dispersed at many locations around the genome, as in *M. bicuspidata* (35). The distribution of empirical allele frequencies from single nucleotide variants found after mapping of Illumina data to the PacBio assembly indicated that the wildtype strain is haploid (36), in contrast to the other reported genome sequences of strains from the *M. pulcherrima/M. fructicola* subclade that are highly heterozygous and probably interspecies hybrids (37, 38). Gene counts for different clusters of orthologous group (COG) categories for the *S. cerevisiae* S288C and *M. pulcherrima* APC 1.2 genomes, analyzed by using the Integrated Microbial Genomes (IMG) system (39), were comparable (Supplementary Figure S5). However, for several categories (i.e., “translation, ribosomal structure and biogenesis”, “replication, recombination and repair”, “posttranslational modification, protein turnover, chaperones”, and “intracellular trafficking, secretion, and vesicular transport”) we counted more genes in *S. cerevisiae*, while the “amino acid transport and metabolism”, “secondary metabolite metabolism”, and “defense mechanisms” categories comprised more genes in the *M. pulcherrima* genome. In order to identify the genetic mutations in the pigmentless *M. pulcherrima* mutant, Illumina HiSeq reads of the three pigmentless mutants W8, W10, and W11 (originating from three individual colonies after plating the same batch of freeze-dried *M. pulcherrima* cells that was stored at 22 °C for several months) were generated and mapped to our *M. pulcherrima* reference genome. The only mutation that was identically detected in all three mutants was a C → A point mutation leading to a premature stop codon at position 1262 of the Mpul 0C08850 gene (Supplementary Table S2). This gene encodes an ortholog of *S. cerevisiae SNF2* and is thus hereafter named Mpul *SNF2* (Figure 3A). Most noteworthy was a point mutation predicted to result in a truncated *M. pulcherrima* Snf2 protein of 420 amino acids in length, as compared to 1578 amino acids for the full-length protein, that lacked all domains and motifs except for the first glutamine-leucine-glutamine (QLQ) domain (Supplementary Figure S6). The functionally relevant helicase and bromo domains (40) were thus not present in the Snf2 protein of the *M. pulcherrima* mutants tested here, which likely resulted in a non-functional Snf2 protein.

**Figure 3:**
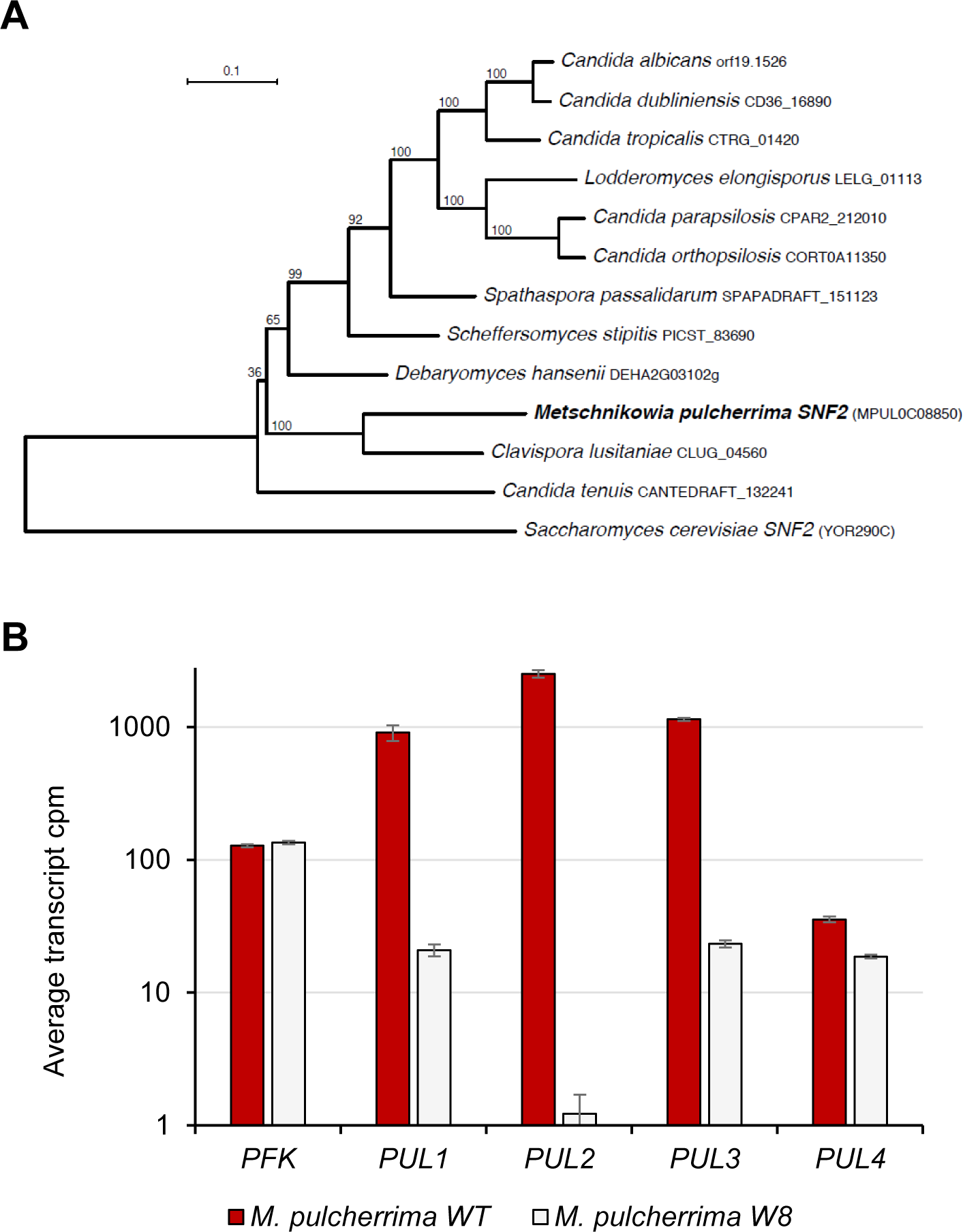
Phylogeny of the *M. pulcherrima* Snf2 protein and *PUL* gene transcription in wildtype and *snf2* mutants. **A)** Phylogenetic relationship of *M. pulcherrima* Snf2 protein to its orthologs in other CUG-Ser clade species, with *S. cerevisiae* Snf2 as outgroup. The tree was constructed by maximum likelihood, and bootstrap supports for internal branches are shown. **B)** The *M. pulcherrima* wildtype (red) and W8 snf2 mutant cells (white) exhibited strongly reduced transcript counts for *PUL1* (Mpul 0C04990), *PUL2* (Mpul 0C04980), and *PUL3* (Mpul 0C04960), while the *PUL4* gene (Mpul 0C04970), encoding a putative transcriptional regulator of the *PUL* genes, was generally lowly transcribed and less affected by the snf2 mutation. As a reference, transcript counts for the 6-phosphofructo-2-kinase (Mpul_0A12740) are shown. The values represent mean transcript counts and the standard error of two and three samples for the wildtype and the mutant, respectively.

Since Snf2 acts as a transcriptional regulator in *S. cerevisiae*, we hypothesized that the *snf2* mutation in the pigmentless *M. pulcherrima* mutants may interfere with transcription of the clustered *PUL* genes that were recently identified to confer pulcherrimin biosynthesis and utilization in *K. lactis* (26). RNAseq data from *M. pulcherrima* wildtype and W8 *snf2* mutant cells confirmed strongly reduced transcript counts for *PUL1* (Mpul 0C04990), *PUL2* (Mpul 0C04980), and *PUL3* (Mpul 0C04960), while the *PUL4* gene (Mpul 0C04970), encoding a putative transcriptional regulator of the *PUL* genes, was generally lowly transcribed and less affected by the *snf2* mutation (Figure 3B). This suggested that lack of pigmentation in *snf2* mutant cells is likely due to a regulatory effect on *PUL* gene transcription.

In summary, we *de novo* assembled a high-quality reference genome for *M. pulcherrima* that served as the basis for identifying a single point mutation in pigmentless, white mutants exhibiting impaired antagonistic activity. An identical Mpul *SNF2* point mutation was found in three pigmentless strains. The point mutation introduced a premature stop codon in *snf2* and likely caused the complex phenotype consisting of reduced antifungal activity and lack of cyclodipeptide synthesis by a transcriptional effect on *PUL* genes and other yet unidentified genes involved in the suppressive activity of *M. pulcherrima*.

### Complementation of pigmentless *M. pulcherrima* mutants with *SNF2* restores antagonistic activity and cyclodipeptide synthesis

To test in an independent experiment whether the premature stop codon in the mutant *snf2* gene was responsible for the lack of pulcherrimin production and the reduced antagonistic activity, a transformation protocol for *M. pulcherrima* was established and the *M. pulcherrima* wildtype *SNF2* gene was reintroduced into the W8 mutant strain by random integration into the genome (Supplementary Figure S7).

Transformation of W8 with a construct containing Mpul *SNF2* under its native promoter, along with a nourseothricin resistance gene selection marker, resulted in pigmented colonies, while transformation with the *snf2* allele did not recover the red phenotype (Supplementary Figure S7). Among 30 transformants growing on the nourseothricin plates, 23 strains exhibited a stable, red phenotype, five strains were unstable and showed red and white colonies after several subculturing rounds, and two strains showed the original white phenotype (Figure 4). Almost all complemented, red strains restored antifungal activity against *G. fujikuroi* to comparable levels as observed for the wildtype (Figure 4). The complemented strain #12 was analyzed in more detail by growth experiments, competition assays against plant pathogenic fungi, bioassays on cherries, and UPLC HR HESI-MS analysis of the metabolites secreted during growth in PDB.

**Figure 4:**
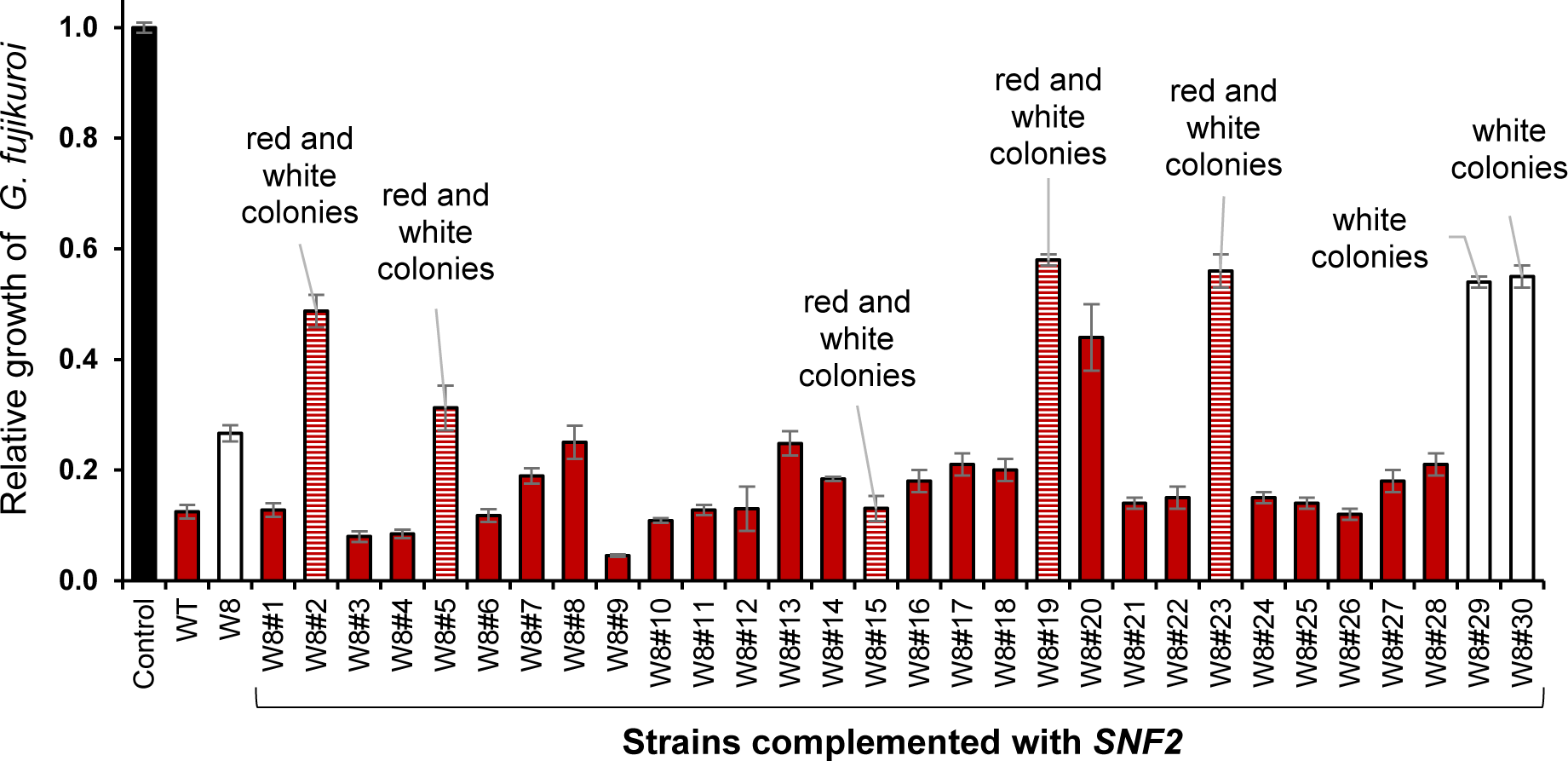
The majority of the complemented strains exhibited a stable, red phenotype and restored antifungal activity. 30 complemented *M. pulcherrima* strains, the original wildtype (APC 1.2) and the pigmentless mutant W8 were tested in binary competition assays against the plant pathogenic fungus *G. fujikuroi*. Growth of *G. fujikuroi* in the presence of the different complemented strains is shown relative to growth in the absence of yeasts (control bar). The color of the bars indicates pigmentation of the wildtype and the complemented strains. In all cases, the mean relative growth of four replicates and the standard error are shown.

Pigmentation of the complemented *M. pulcherrima* strain W8#12 on agar plates with increasing iron concentrations was similar to the wildtype and remained stable (Figure 1A). The antagonistic activity was also restored to wildtype levels, as indicated by binary competition assays on agar plates against *B. caroliniana, G. fujikuroi*, and *F. oxysporum* isolates (Figures 1B, 1C; Supplementary Figure S1A). Likewise, on cherries, the complemented *M. pulcherrima* mutant W8#12 showed restored antagonistic activity both with respect to rot diameter and mycelium development in artificially wounded and infected cherry fruits (Figures 1D, 1E). Finally, introduction of the wildtype *SNF2* gene restored the secretion of **1**, cyclo(L-leucyl-L-leucyl), **2**, pulcherriminic acid, as well as an unidentified precursor **3** and two probable degradation products **4** and **5,** into the culture supernatants of the complemented strain #12 (Figure 2C). Overall, these results defined a point mutation in Mpul *SNF2* as the underlying cause for abolished pigmentation, lack of pulcherriminic acid secretion and reduced antagonistic and biocontrol activity against plant pathogenic fungi.

## Discussion

*Metschnikowia pulcherrima* is a strongly antifungal yeast that is frequently identified in the phyllosphere and considered a promising species for biocontrol applications. Its most prominent characteristic is the formation of a red pigment, pulcherrimin, that was discovered over 60 years ago (22). This compound represents an insoluble complex of iron and pulcherriminic acid that forms non-enzymatically in the presence of iron III (22, 24, 25, 41, 42). Pulcherrimin formation may serve as a defence against deleterious effects of high amounts of iron (reviewed by Kluyver (22)) and is thought to deprive competing microbes of iron and thus limit their growth (9, 16, 30, 32, 33). Although pulcherrimin levels and antifungal activity seem to correlate, as suggested previously (33) and shown here by quantitative competition assays with pigmentless and complemented mutants, a causal relationship between iron deprivation and antifungal activity has not been demonstrated. The function of pulcherrimin has thus not been elucidated in detail yet and the role of iron and iron chelation is not resolved. For example, pulcherriminic acid is also synthesised under iron replete conditions, which is the reason why antimicrobial activities irrespective of iron or iron monopolisation, rather than iron scavenging, have also been hypothesised as modes of action (26, 33, 43). It has also been observed that the addition of iron may, in some cases, increase or at least not significantly diminish antifungal activity (7, 9), which would not be expected if iron depletion was the mode of action. While a recent study identified a putative transporter enabling yeast to take up and utilise pulcherrimin as a proper siderophore (26), it has also been doubted whether or not iron can be taken up from the insoluble pulcherrimin complex and serve as an iron source for the synthesising organisms (e.g., bacteria and yeasts) (30). In *Bacillus subtilis,* where pulcherrimin biosynthesis has been studied in more detail, the formation of the pulcherrimin precursor pulcherriminic acid does not depend on iron availability and the main repressor of pulcherriminic acid biosynthesis (YvmB) is even upregulated under iron starvation (31, 43), also arguing against a role of pulcherrimin in iron provision. It has therefore been suggested that iron chelation by pulcherriminic acid may interfere with iron-dependent cellular process in general and thereby indirectly cause diverse changes in the cell (31).

The study presented here was initiated by the discovery of spontaneously occurring, pigmentless *M. pulcherrima* mutants. The pigmentless phenotype was regularly observed after plating freeze dried cells that had been stored at 22 °C for several months; at a time point when the viability of the stored cells had already strongly diminished. Unexpectedly, the three pigmentless strains, originating from three independent, pigmentless colonies, harboured the same *snf2* point mutation. Since it seems unlikely that the same point mutation occurred three times independently in the freeze-dried cells, this genetic alteration has likely arisen already during fermentation. It is also possible that the pigmentless phenotype confers a selective advantage for long-term storage in the freeze-dried powder at elevated temperature. The competitiveness and sensitivity to stress conditions, such as long-term storage, of the mutant and wildtype *M. pulcherrima* cells will have to be studied in detail to test this hypothesis. Under the standard growth and assay conditions used throughout the studies shown here, the pigmentless *M. pulcherrima* mutants did not exhibit any growth defect; if anything, the mutants grew slightly better, as could be expected based on the lack of an antimicrobial metabolite. The red pigment pulcherrimin has caught the interest of research already more than 60 years ago, but much about its chemical nature, synthesis and biological function remains to be elucidated. Here, we formally prove the presence of cyclo(L-leucyl-L-leucyl) and four congeners, including the pulcherrimin precursor pulcherriminic acid. UPLC HR HESI-MS in combination with labelling and feeding experiments provided not only exact molecular formulas but also biosynthetic information that allowed us to identify two possible degradation products of pulcherriminc acid or pulcherrimin. These products may arise from enzymatic reactions involving monooxygenation and decarboxylation, but the exact mechanism still has to be elucidated. In addition, it is not yet clear if these new intermediates and possible degradation products contribute to the overall antifungal activity of *M. pulcherrima* and what the biological function of pulcherriminic acid/pulcherrimin degradation is.

Although the reduced antifungal activity of the pigmentless *M. pulcherrima* cells supports the notion that pulcherrimin may indeed be responsible for the suppressive phenotype, it is important to note that pigmentless mutants only showed reduced antifungal activity and still strongly inhibited the growth of filamentous fungi. These results clearly demonstrate that the *M. pulcherrima* antifungal activity depends on several mechanisms of which pulcherrimin may be one. The nature and interplay of the mechanisms contributing to *M. pulcherrima* antifungal activity, including pulcherrimin formation, are thus still largely unknown and need to be studied and elucidated in much more detail.

The mutation identified to cause the pigmentless phenotype in *M. pulcherrima* affected the homolog of the *S. cerevisiae SNF2* gene; a point mutation caused a premature stop codon and thus encoded for a truncated Mpul Snf2 protein. In *S. cerevisiae*, Snf2 is a non-essential ATPase that modulates nucleosome structure as part of the SWI/SNF chromatin remodelling complex and thus regulates transcription of a plethora of genes (44-46). The gene/protein name is derived from the sucrose non-fermenting phenotype (47, 48), but *SNF2* deletion and mutations strongly affect yeast metabolism overall (49) and cause a diverse range of phenotypes including abolished mating type switching and sporulation, decreased resistance (e.g., to chemicals, metals, killer toxin), or altered cytosolic pH (50-55). In humans, mutations in the *SNF2* homolog, and components of the SWI/SNF complex in general, are frequently identified in different types of cancers, leukemia, and other disorders (56-59). In this study, a point mutation leading to a premature stop codon in *SNF2* caused a pigmentless phenotype by abolishing cyclodipeptide synthesis (i.e., synthesis of cyclo(Leu-Leu) and pulcherriminic acid) and reduced antifungal activity as demonstrated by competition assays on plates and bioassays on fruits against plant pathogenic fungi. Our results thus link Snf2 activity to the regulation of pulcherrimin metabolism and antifungal activity. This was confirmed by the strongly reduced transcript levels of the recently identified pulcherrimin biosynthesis genes *PUL1* (Mpul 0C04990) and *PUL2* (Mpul 0C04980) in the *snf* mutant strain as compared to the wildtype.

In summary, this multidisciplinary study identified a point mutation in the *M. pulcherrima SNF2* gene as the underlying cause for lack of pigmentation, abolished cyclodipeptide biosynthesis, and reduced antifungal activity. This identification of Snf2 as the first “biocontrol regulator” thus serves as a starting point and enabler for further elucidating the regulation and underlying mechanisms of pulcherrimin biosynthesis, function and metabolism in *M. pulcherrima*, which will hopefully benefit biocontrol applications of this yeast in the future.

## Supporting information

## Experimental procedures

### Isolates and cultivation

The wildtype *M. pulcherrima* (Pitt & M.W. Mill.) (SH180747.07FU; isolate APC 1.2, CCOS978 (Culture Collection of Switzerland)) was originally isolated from apple flowers in Switzerland (4). For competition assays, the filamentous fungi *Botrytis caroliniana* X.P. Li & Schnabel (isolate EC 1.05, SH177344.07FU), *Gibberella fujikuroi* ((Sawada) Wollenw.) (SH213620.07FU; isolate BC 8.14, CCOS1020), and *Fusarium oxysporum* f. sp. *radicis-lycopersici* (Schlecht as emended by Snyder and Hansen) (NRRL 26381/CL57 (ARS Culture Collection, USDA)) were used. All isolates were identified by sequencing the ITS region and assigning a UNITE species hypothesis (4, 60). The pigmentless mutants W8, W10 and W11 were isolated from cells that were plated out after extended storage as freeze-dried powder at 22°C. All *M. pulcherrima* strains were maintained on potato dextrose agar (PDA; Becton, Dickinson and Company, Le Pont de Claix, France or Formedium™, Norfolk, United Kingdom) plates, grown at 22°C, and transferred to fresh plates weekly.

### Competition assays

The protocol for binary competition assays was based on the procedure by Hilber-Bodmer et al. (4), but included minor adjustments. Yeasts cells and conidia of filamentous fungi were collected in water and adjusted to an OD_600_ of 0.001 and 0.1, respectively. 15 or 30 μl of the yeast suspension was plated on PDA plates of 5.5 or 9 cm diameter, respectively, and 5 μl of the conidial suspension were inoculated in the center of the plates. Plates were incubated at 22°C for 3-15 days depending on the fungal species. Growth of the filamentous fungus was quantified before it reached the edge of the control plate (plate without yeasts) with the help of a planimeter (Planix 5, Tamaya Technics Inc., Tokyo, Japan). The average of the relative growth (growth in presence of yeast/growth on control plate) of four replicates was calculated. All assays were repeated at least twice and showed comparable results.

For bioassays, conventionally grown cherries (1-2 weeks in cold storage) were washed with 70% ethanol and water, and a 2.5 mm wide and 3 mm deep lesion was created at the equator of each fruit by using a custom-made tool. After complete drying of the lesion, 10 μl of a *M. pulcherrima* suspension (OD_600_ of 1) or sterile water as a control was applied to the lesion. After 3 h, 10 μl of a *Botrytis* conidia suspensions (10^5^ conidia/ml) or water as a control was added to the lesions. The cherries of all treatments were kept on a dry filter paper in plastic trays, with a wet filter paper (soaked with 20 ml water) under the trays, and stored within a plastic bag at 22°C. After 4 days, rot and mycelium diameter were measured. For each treatment, 15 cherries were used and the experiment was performed twice for each treatment. The mean and standard errors of one representative experiment are shown.

### Identification of secreted cyclic dipeptides

20 ml PDB containing 400 μl Amberlite XAD16N slurry (Sigma-Aldrich Chemie GmbH, Buchs, Switzerland) were inoculated with *M. pulcherrima* cells (wildtype, mutant, or complemented cells) to an OD_600_ of 0.1 and grown at 22°C for 2-3 d. For labelling experiments, yeast nitrogen base medium (without amino acids; Formedium™, Norfolk, United Kingdom) was prepared with ^13^C_6_ D-Glucose (Cambridge Isotope Laboratories / ReseaChem GmbH, Burgdorf, Switzerland) and ammonium sulfate or with glucose and ^15^N_2_ ammonium sulfate (Sigma-Aldrich Chemie GmbH, Buchs, Switzerland). The growth medium was discarded, the XAD beads were washed three times with water, and eluted with 1 ml methanol. The solvent was evaporated in a SpeedVac, the extracted material dissolved in 1 ml water, and the sample stored at −20°C until further analysis. All extracts and a cyclo(Leu-Leu) standard (ChemFaces Biochemical Co., Ltd., Wuhan, China) were analyzed by ultra-performance liquid chromatography-high resolution heated electrospray ionization mass spectrometry (UPLC HR HESI-MS). Data were recorded on a Thermo Scientific™ Q Exactive™ Hybrid Quadrupole-Orbitrap Mass Spectrometer coupled to a Dionex Ultimate 3000 UPLC. We used the following solvent gradient (A = H_2_0 + 0.1% formic acid, B = acetonitrile + 0.1% formic acid with B 5-20% from 0-2 min, 20-98% from 2-20 min, 98% from 20-25 min, 98-5% from 25-27 min at a flowrate of 0.5 ml/min) on a Phenomenex Kinetex 2.6 μm XB-C18 100 Å (150 × 4.6 mm) column at 30°C. The MS was operated in positive ionization mode at a scan range of 200-2000 *m*/*z* and a resolution of 70000. The spray voltage was set to 3.5 kV, the S-lens to 50, the auxiliary gas heater temperature to 438°C and the capillary temperature to 270°C. Further parameters used were AGC target (1e6), maximum injection time (200 ms), microscans (1), sheath gas (53), aux gas (14) and sweep gas (3).

### Genome sequencing & mapping of mutations

Genomic DNA of the *M. pulcherrima* strain APC 1.2 was extracted using the Qiagen DNeasy Plant Mini Kit and sequenced on the PacBio RS II platform (performed at the Functional Genomics Center Zurich). Subsequent *de novo* genome assembly, polishing and resequencing were performed using PacBio SMRT Portal 2.3.0 (61). The genomes of three mutant *M. pulcherrima* colonies (W8, W10 and W11) were sequenced using the Illumina MiSeq technology (Micro flow cell, paired-end, 250 bp), producing 1’225’187 W8, 1’087’938 W10 and 1’054’343 W11 reads. They were mapped to the *M. pulcherrima* reference genome using the BWA-MEM algorithm version 0.7.15. (62). Secondary and supplementary alignments were removed subsequently using SAMtools (version 1.3.1) (63).

### Genome annotation, phylogenetic analysis, and functional characterisation

Genome annotation was performed using a modified version of the Yeast Genome Annotation Pipeline (YGAP; (64)), which enabled to use the homology and synteny information from the Candida Gene Order Browser database instead of the Yeast Gene Order Browser database (CGOB and YGOB; (65, 66)). The CGOB database is composed of 13 species within the *Candida* (CUG-Ser1) clade, which are phylogenetically closer to *M. pulcherrima* than are the species in YGOB. YGAP was therefore modified to account for the alternative genetic code and to use a different ancestral reference for the synteny-based annotation. Because an ancestral gene order has not been yet inferred for the species in CGOB, the *Clavispora lusitanae* genome (67) was used for synteny reference as it was found to be the closest relative to *M. pulcherrima* among the species in CGOB. The phylogenetic tree in Figure 3A was constructed from an alignment of orthologous Snf2 proteins taken from CGOB, using MUSCLE, Gblocks and PhyML (1013 sites, GTR+gamma substitution model, 4 rate categories, 100 bootstrap replicates) as implemented in SeaView v4.50 (68) with their default parameters. The annotated genome of the wildtype *M. pulcherrima* strain APC 1.2 was functionally analyzed and is being distributed by the Integrated Microbial Genomes (IMG) system of the Department of Energy’s (DOE’s) Joint Genome Institute (JGI) (IMG genome ID 2721755843; https://genome.jgi.doe.gov/portal/IMG_2721755843/IMG_2721755843.info.html) (39). The *M. pulcherrima* APC 1.2 genome is also available at NCBI under the BioProject PRJNA508581 (Accessions CP034456-CP034462 for the seven scaffolds).

### RNAseq experiment

Wildtype and mutant (W8) *M. pulcherrima* cells were grown in potato dextrose broth (PDB; Formedium™, Norfolk, United Kingdom). An overnight culture was diluted with fresh medium to an OD_600_ of 0.4. After 5-6 h of growth at 23°C in a shaking incubator, 1 OD_600_ was collected, spun down, the supernatant was discarded, and the pellet frozen in liquid nitrogen and stored for later RNA extraction. Cells were mechanically disrupted with a FastPrep FP120 (speed 4.5, time 37 s; Thermo ELECTRON Corporation) and total RNA was extracted with the Qiagen RNeasy mini kit according to the manufacturers protocol and submitted for RNA sequencing to the Functional Genomics Center Zurich. RNA was quantified and verified with a Qubit^®^ 3 fluorometer (Invitrogen) and a Bioanalyzer 2100 (Agilent, Waldbronn, Germany). The TruSeq Stranded mRNA Sample Prep Kit (Illumina, Inc, California, USA) was used for RNA depletion and reverse transcription of 1 μg of total RNA samples. The cDNA samples were fragmented, end-repaired and polyadenylated before ligation of TruSeq adapters for multiplexing. Fragments containing TruSeq adapters on both ends were selectively enriched with PCR. The quality and quantity of the enriched libraries were validated using Qubit^®^ (1.0) Fluorometer and the Bioanalyzer 2100 (Agilent, Waldbronn, Germany). The TruSeq SR Cluster Kit v4-cBot-HS or TruSeq PE Cluster Kit v4-cBot-HS (Illumina, Inc, California, USA) was used for cluster generation using 8 pM of pooled normalized libraries on the cBOT. Sequencing was performed on the Illumina HiSeq 2500 paired end at 2 X126 bp or single end 126 bp using the TruSeq SBS Kit v4-HS (Illumina, Inc, California, USA). For the wildtype and the W8 mutant strain, two and three independent biological replicates were used for RNA extraction and consecutive sequencing, respectively. The RNAseq data were analyzed with an early version of the workflow available at https://github.com/csoneson/rnaseqworkflow. Briefly, reads were trimmed with TrimGalore! (https://www.bioinformatics.babraham.ac.uk/projects/trim_galore/) to remove adapters and low-quality sequences. Gene expression levels were estimated using Salmon (69). To identify differentially expressed genes and estimate normalized expression values (counts per million), the counts were subsequently analyzed with edgeR (69). The RNAseq data have been deposited in NCBI’s Gene Expression Omnibus (70) and are accessible through GEO Series accession number GSExxx (https://www.ncbi.nlm.nih.gov/geo/query/acc.cgi?acc=GSExxx) *(submission is being processed)*.

### Complementation of mutants

Wildtype and mutant *SNF2* coding sequences along with the flanking promoter and terminator sequences were PCR-amplified using Q5 High-fidelity polymerase (NEB) and primers 5’-caactagtAGTCAGCGTCACTTAGTGTGAAATACCTG and 5’-catgtcgacGTCCATCAATCGAATCAGCATAATACTGCC. Cycling conditions were as per manufacturer’s instructions, with an annealing temperature of 64°C and an extension time of 3 minutes 15 seconds. Purified PCR products were ligated (T4 DNA ligase, NEB) via SpeI and SalI (NEB) restriction sites into a pEX-A2 plasmid backbone containing a codon-optimised nourseothricin (Nat, Werner Bioagents) yeast selection marker under the strong constitutive TEF promoter previously cloned from the *M. pulcherrima* genome. Ligations were used to transform competent *E.coli* (NEB, 5-alpha) with selection by ampicillin (Sigma, 100μg/ml) and Nat (50μg/ml). Positive colonies identified via colony PCR (DreamTaq Green PCR Mastermix, ThermoFisher; primers 5’-GCATGATGTGACTGTCGCCCGTAC and 5’-CCGTACTGGGCCTTGAGCTG; cycling conditions as per the manufacturer’s instructions, with an annealing temperature of 59°C and an extension time of 1 minute) were propagated overnight in Luria broth (Sigma) supplemented with 100μg/ml ampicillin (Sigma) and 50μg/ml Nat and plasmids purified via miniprep. All protocols used in the above cloning steps were as per the manufacturer’s instructions and all purification steps performed using GeneJET kits (ThermoFisher).

Plasmids containing the wildtype or mutant gene clone were digested with Acc65I and SalIHF (NEB) and used to transform both the WT and W8 *M. pulcherrima* strains using a lithium acetate method (71). In brief, overnight cultures of the yeast strains were diluted to a starting OD_600_ of 0.3 and allowed to grow to an OD_600_ of 0.9. 1ml of culture per transformation was centrifuged, the cell pellet washed once with PBS then resuspended in a transformation mix comprised of 800μl 50% PEG-3350 + 100μl 1M LiOAc pH 7.4 + 40μl 5mg/ml boiled salmon sperm + 100μl 10XTE + 20μl 1M DTT + 1μg linearised plasmid. Transformations were incubated overnight (25°C, static), heat-shocked at 40°C for 5 minutes, placed on ice for one minute then centrifuged for 5 minutes at 4000rpm. The supernatant was removed, cells resuspended in 600μl of SMB (30g/l Tryptic Soy Broth (Sigma), 25g/L Malt Extract (Sigma), pH5) and incubated for 1 hour (25°C, 200rpm) before plating on to Malt Extract Agar (MEA, Sigma) plates containing 50μg/ml Nat. After two days at 25°C, yeast colonies that grew were patched on to a second MEA+Nat plate. Colonies that grew overnight at 25°C were then subject to colony PCR to confirm the presence of the plasmid (primers and conditions as per colony PCR to confirm positive *E.coli* colonies above) and scored for colour (red or white).

## Acknowledgements

Maja Hilber-Bodmer, Liesa Kunz, Chloé Douard, Andrea Knauf and Denise Müller are acknowledged for help with experimental work. We are grateful to Prof. Mark Robinson (University of Zurich) and Dr. Charlotte Soneson (Friedrich Miescher Institute for Biomedical Research, Basel) for access to an early version of their RNASeq data analysis workflow. Michael Schmid and Vincent Sommerville helped with genome assembly and RNAseq analysis, respectively. The project is supported by the Swiss National Science Foundation (SNSF, grant 31003A_175665 / 1) to FMF. DGL and MMB are funded by the Industrial Biotechnology Catalyst (Innovate UK, BBSRC, EPSRC) to support the translation, development and commercialisation of innovative Industrial Biotechnology processes (EP/N013522/1).

## Author contributions

Responsibilities for the different methods and sections of the manuscript was shared as follows: DGL, MMB, and DH established the transformation protocol for *M. pulcherrima* and generated the complemented strains. KHW and RAOM annotated and analyzed the *M. pulcherrima* genome and *SNF2* gene. KS and CHA sequenced and assembled the *M. pulcherrima* genome and identified the *snf2* point mutation. AB and JP performed the UPLC HR HESI-MS analyses and thus identified cyclo(Leu-Leu), pulcherriminic acid and the precursors and degradation products of these compounds. IS, MS, PK, ASK performed all competition assays, growth experiments, and grew the cultures for DNA, RNA and metabolite analyses. FMF isolated the pigmentless mutants, analyzed *PUL* gene transcription, coordinated the study, and wrote the majority of the manuscript. All authors have read and agree with the manuscript.

