## Supplementary material for "Snf2 controls pulcherriminic acid biosynthesis and connects pigmentation and antifungal activity of the yeast *Metschnikowia pulcherrima*"

### Supplementary Tables

**Supplementary Table S1.** Assembly and genome statistics for *M. pulcherrima*, *M. fructicola*, and *M. bicuspidata*. The statistics refer to the *Metschnikowia bicuspidata* NRRL YB-4993 (35) and *M. fructicola* strain 277(38) genomes. For *M. pulcherrima*, the 7 scaffolds refer to the nuclear genome (excluding the mitochondrial genome).

| | ***M. pulcherrima*** | *M. fructicola* | *M. bicuspidata* |
|---|---|---|---|
| technology | **PacBio** | PacBio | 454, Illumina |
| assembler | **Canu** | HGAP 3.0 | Newbler v2.5 |
| coverage (x) | **254** | 20 | 16 |
| assembly (Mbp) | **15.88** | 24.48 | 16.06 |
| contigs (n) | **7** | 93 (93 scaffolds) | 421 (33 scaffolds) |
| $N_{50}$ (bp) | **2’688’338** | 957’836 | 62’344 |
| GC (%) | **46** | 46 | 47 |
| proteins (n) | **5922 (genes)** | NA | 5,851 |
| tRNAs (n) | **227** | NA | NA |

**Supplementary Table S2.** Variant detection in the three pigmentless *M. pulcherrima* mutants W8, W10 and W11. The amino acid changes caused by the mutations are indicated by the amino acid one letter code. “*” indicates a stop codon. Synonymous mutations not leading to a different amino acid sequence are labelled “SNP syn.

| Identified gene | Mutant strain | Mean read coverage | Ref. sequence | Ref. Count | Mutated sequence | Mut. Count | Impact |
|---|---|---|---|---|---|---|---|
| Mpul 0A10550 | W8 | 30.3 | G | 0 | T | 21 | SNP P → Q |
| Mpul 0A02060 | W10 | 27.7 | C | 1 | A | 34 | SNP L → I |
| Mpul 0A02190 | W10 | 27.7 | A | 2 | T | 26 | SNP S → Q |
| Mpul 0A03950 | W10 | 27.7 | C | 0 | T | 27 | SNP syn. |
| Mpul 0A04630 | W10 | 27.7 | A | 0 | G | 31 | SNP syn. |
| Mpul 0A13480 | W10 | 27.7 | GGC | 0 | - | 15 | Triplet deletion |
| Mpul 0B06920 | W8 | 31.6 | G | 1 | A | 23 | SNP V → I |
| Mpul 0B11570 | W10 | 28.7 | G | 3 | T | 37 | SNP L → I |
| **Mpul 0C08850** | **W8** | **31.8** | **C** | **0** | **A** | **33** | **SNP S → *** |
| **Mpul 0C08850** | **W10** | **28.9** | **C** | **0** | **A** | **31** | **SNP S → *** |
| **Mpul 0C08850** | **W11** | **27.6** | **C** | **0** | **A** | **31** | **SNP S → *** |
| Mpul 0D05330 | W10 | 27.6 | T | 1 | C | 29 | SNP syn. |
| Mpul 0F05540 | W8 | 30.9 | A | 0 | G | 24 | SNP D → G |
| Mpul 0G02320 | W8 | 29.7 | - | 0 | C | 11 | Longer ORF |

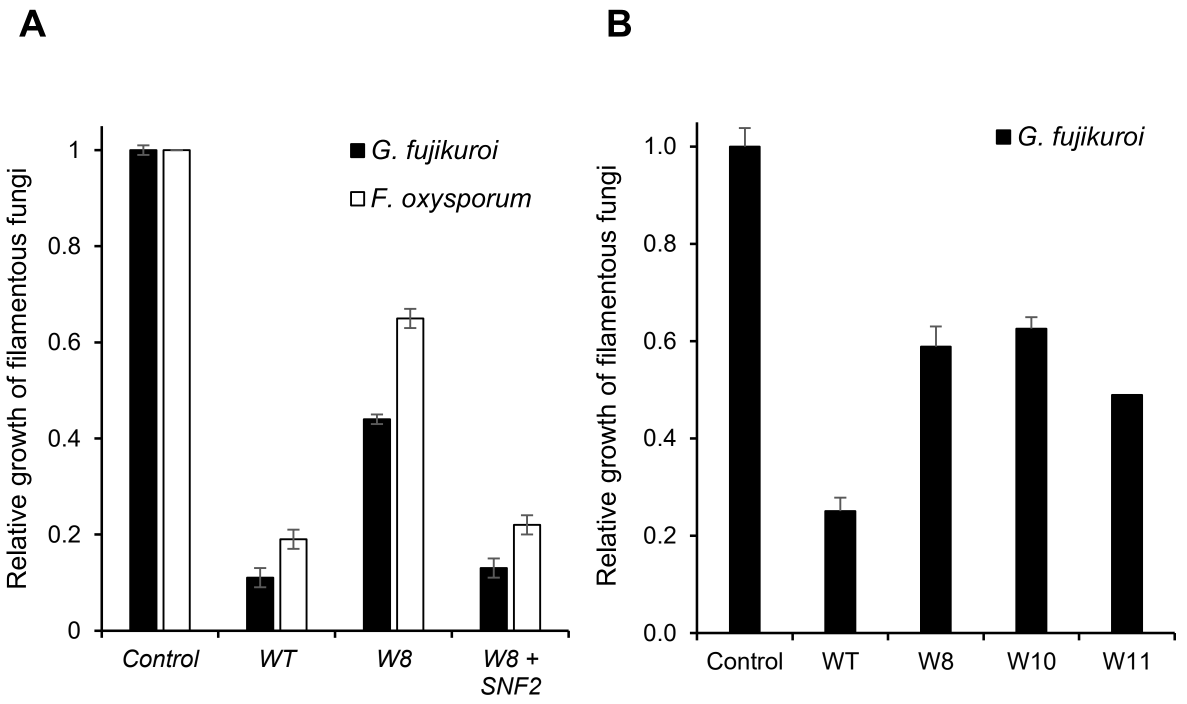


**Supplementary Figure S1**: Pigmentless *M. pulcherrima* mutants are less antagonistic than the wildtype and complementation with SNF2 restores antifungal activity. A) The antifungal activity of the *M. pulcherrima* wildtype (APC 1.2) and W8 mutant was tested against *G. fujikuroi* and *F. oxysporum*. B) The three pigmentless *M. pulcherrima* mutants W8, W10 and W11 were tested in binary competition assays against *G. fujikuroi*. In all cases, the mean and standard error of four quantifications is shown.

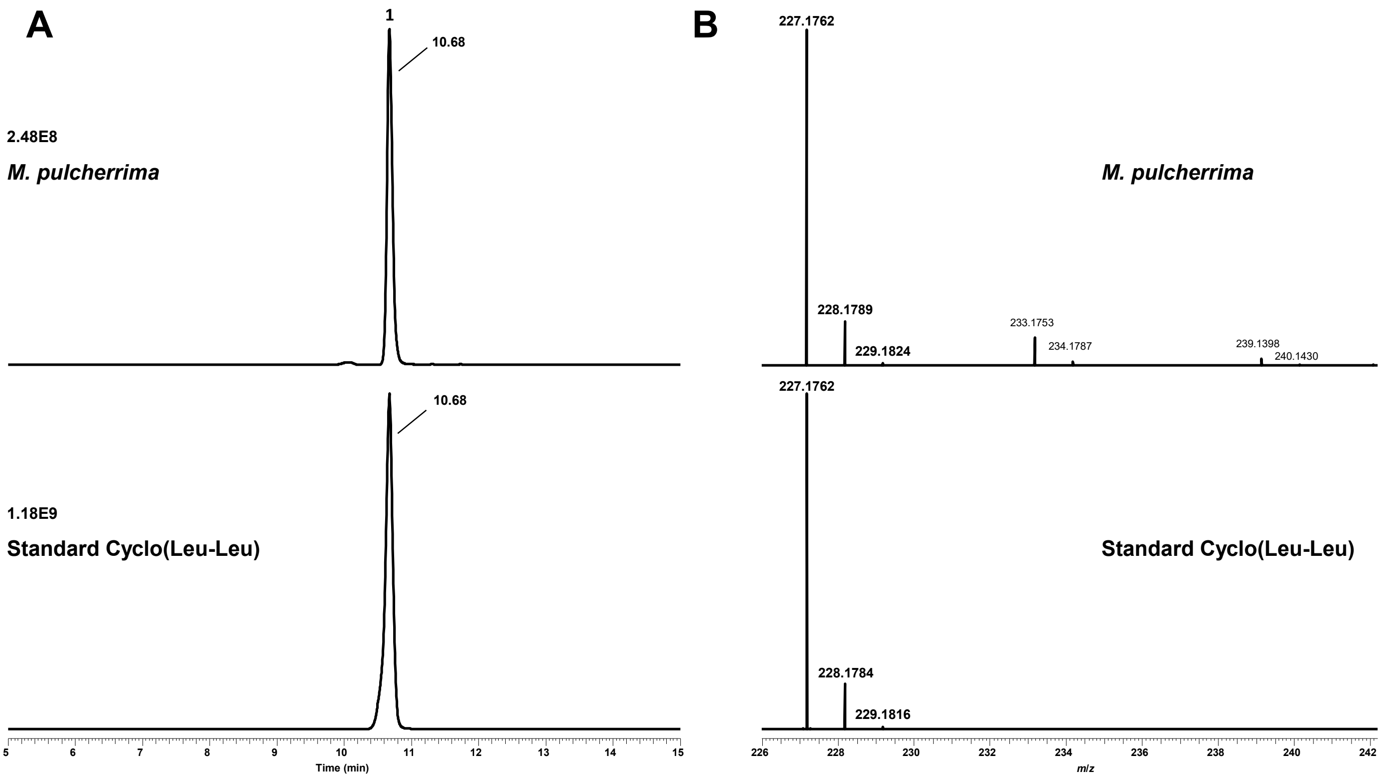


**Supplementary Figure S2**: Comparison of **1** $[M+H]^+$=227.1672 *m*/*z* with synthetic standard Cyclo(Leu-Leu). A) Retention times of **1** and Cyclo(Leu-Leu) B) Mass spectra of **1** and Cyclo(Leu-Leu)

*Supplementary Figure S3*

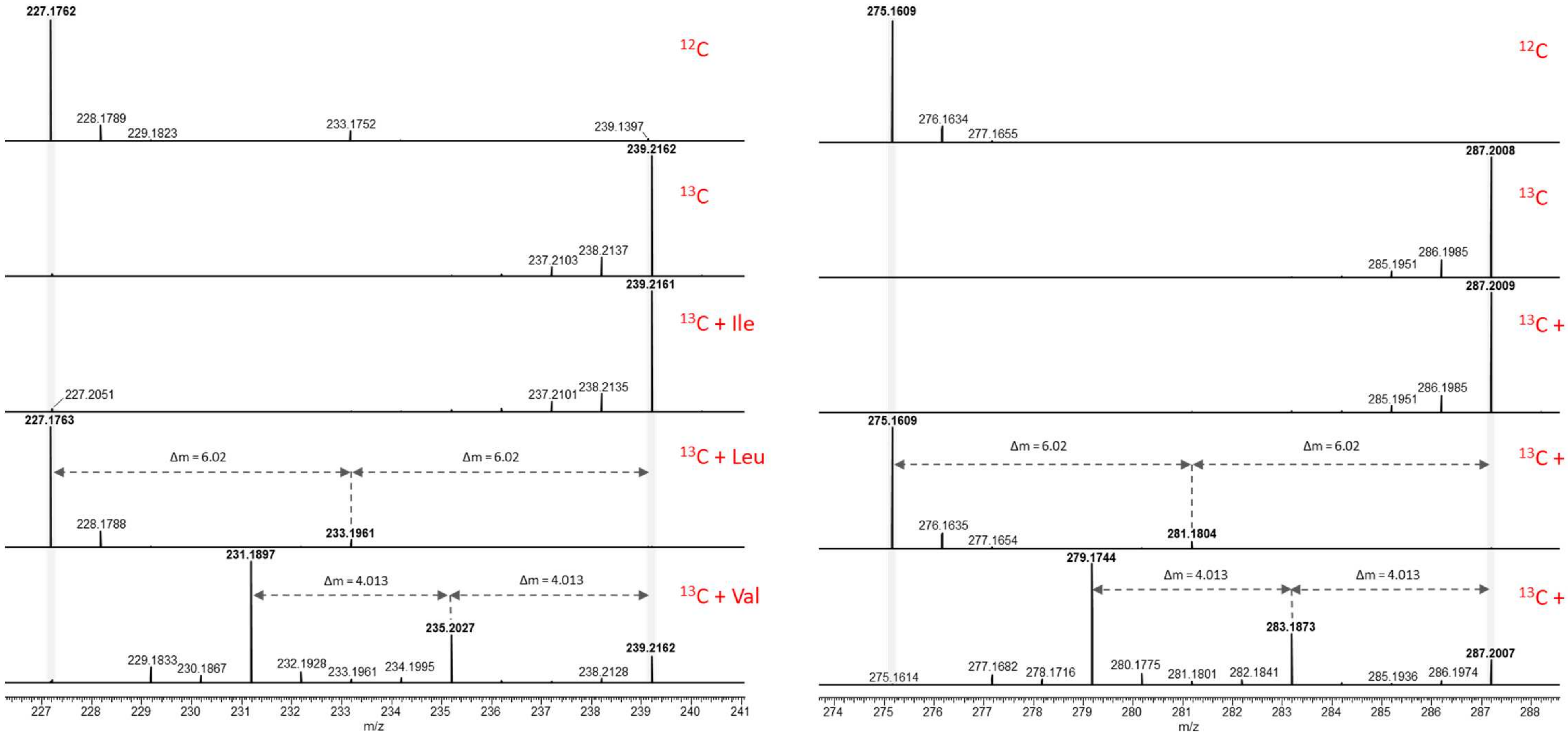


*Mpul SNF2*

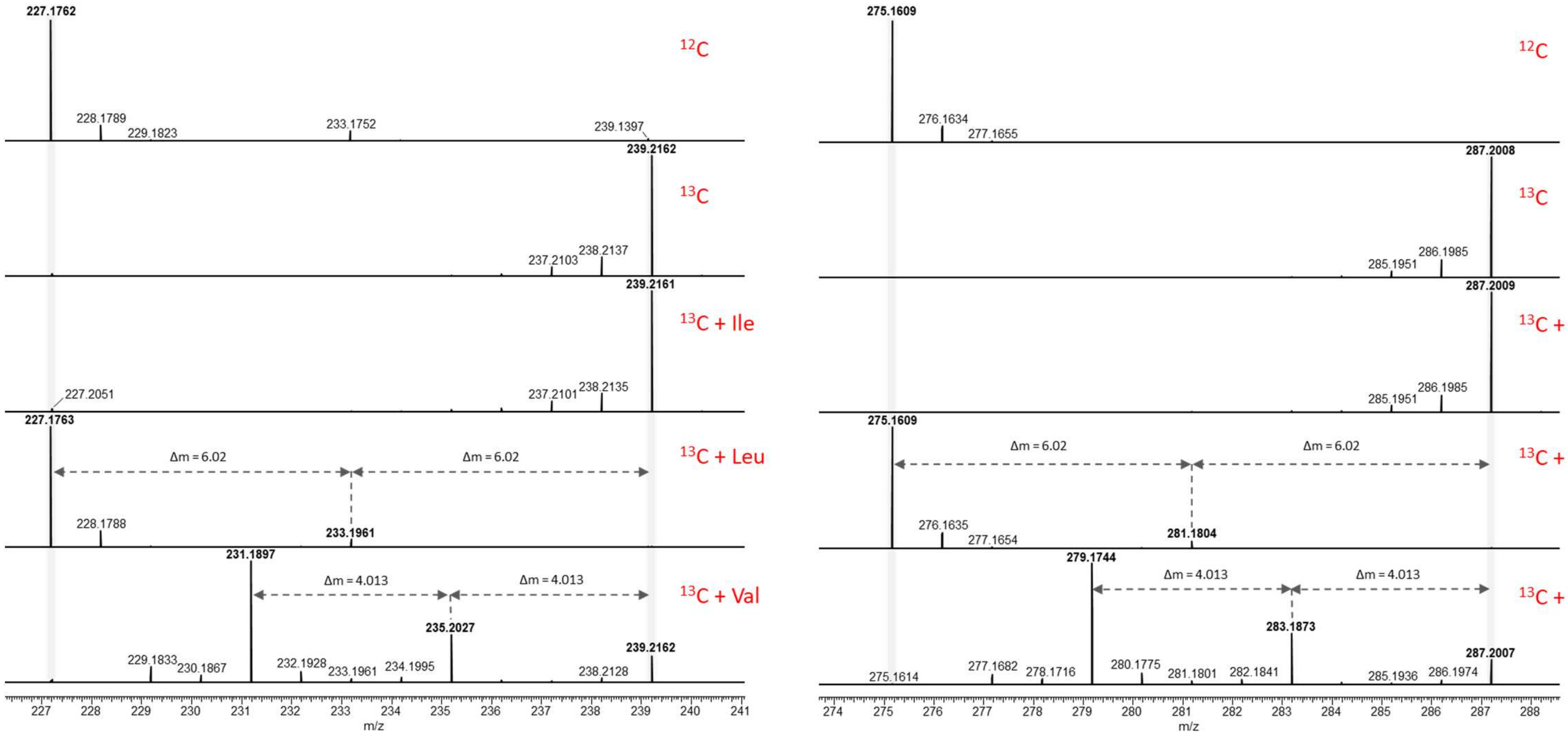

227.1762
228.1789
229.1823
233.1752
239.1397
12C
239.2162
13C
237.2103
238.2137
239.2161
13C + Ile
227.2051
237.2101
238.2135
227.1763
13C + Leu
Δm = 6.02
Δm = 6.02
228.1788
233.1961
231.1897
Δm = 4.013
Δm = 4.013
13C + Val
235.2027
239.2162
229.1833
230.1867
232.1928
233.1961
234.1995
238.2128
227 228 229 230 231 232 233 234 235 236 237 238 239 240 241
m/z
275.1609
276.1634
277.1655
12C
287.2008
13C
285.1951
286.1985
287.2009
13C + Ile
285.1951
286.1985
275.1609
Δm = 6.02
Δm = 6.02
13C + Leu
276.1635
277.1654
281.1804
279.1744
Δm = 4.013
Δm = 4.013
13C + Val
283.1873
287.2007
275.1614
277.1682
278.1716
280.1775
281.1801
282.1841
285.1936
286.1974
274 275 276 277 278 279 280 281 282 283 284 285 286 287 288
m/z


*Mpul SNF2*

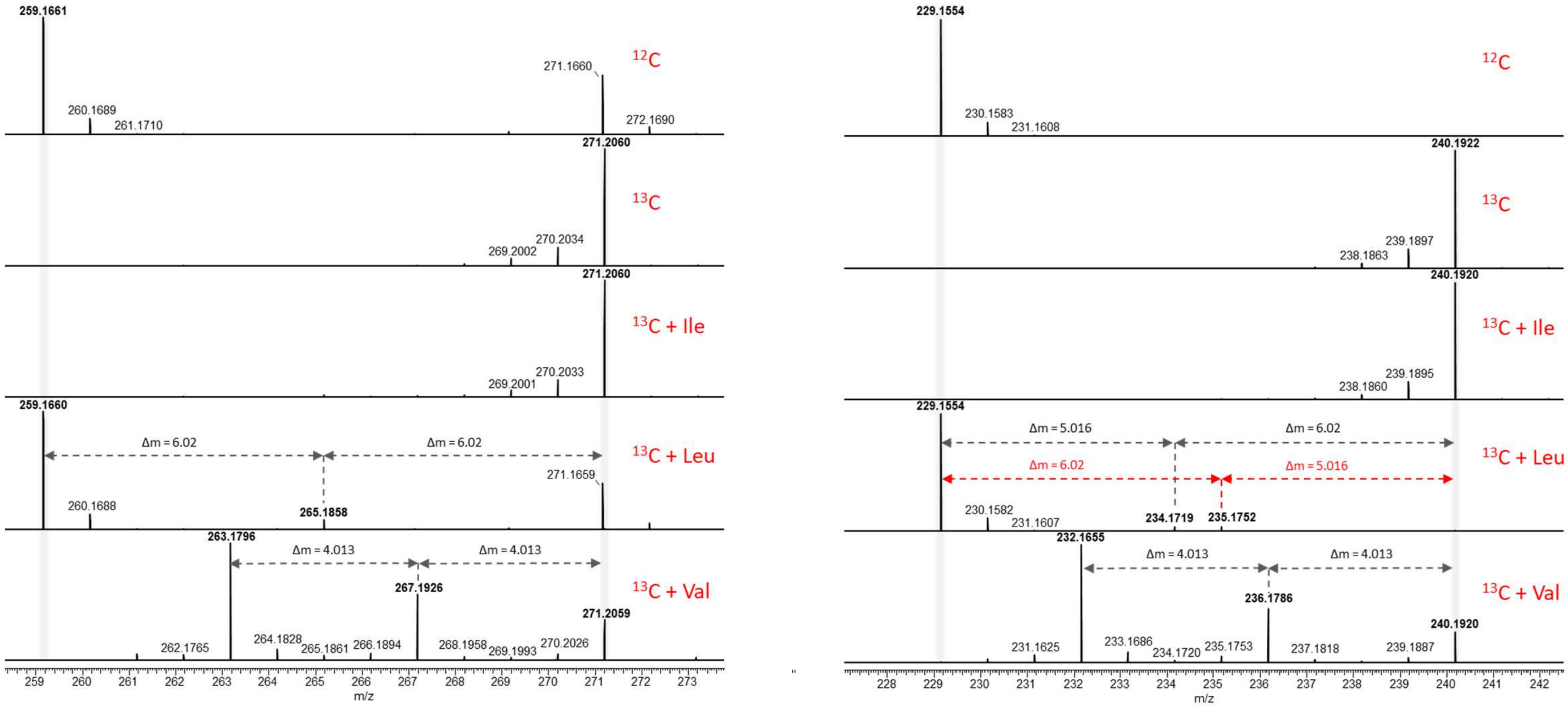
259.1661
260.1689
261.1710
271.1660
272.1690
12C
271.2060
269.2002
270.2034
13C
271.2060
269.2001
270.2033
13C + Ile
259.1660
260.1688
265.1858
271.1659
Δm = 6.02
Δm = 6.02
13C + Leu
263.1796
262.1765
264.1828
265.1861
266.1894
267.1926
268.1958
269.1993
270.2026
271.2059
Δm = 4.013
Δm = 4.013
13C + Val
259 260 261 262 263 264 265 266 267 268 269 270 271 272 273
m/z
229.1554
230.1583
231.1608
12C
240.1922
238.1863
239.1897
13C
240.1920
238.1860
239.1895
13C + Ile
229.1554
230.1582
231.1607
234.1719
235.1752
Δm = 5.016
Δm = 6.02
Δm = 6.02
Δm = 5.016
13C + Leu
232.1655
231.1625
233.1686
234.1720
235.1753
236.1786
237.1818
239.1887
240.1920
Δm = 4.013
Δm = 4.013
13C + Val
228 229 230 231 232 233 234 235 236 237 238 239 240 241 242
m/z

*Mpul SNF2*

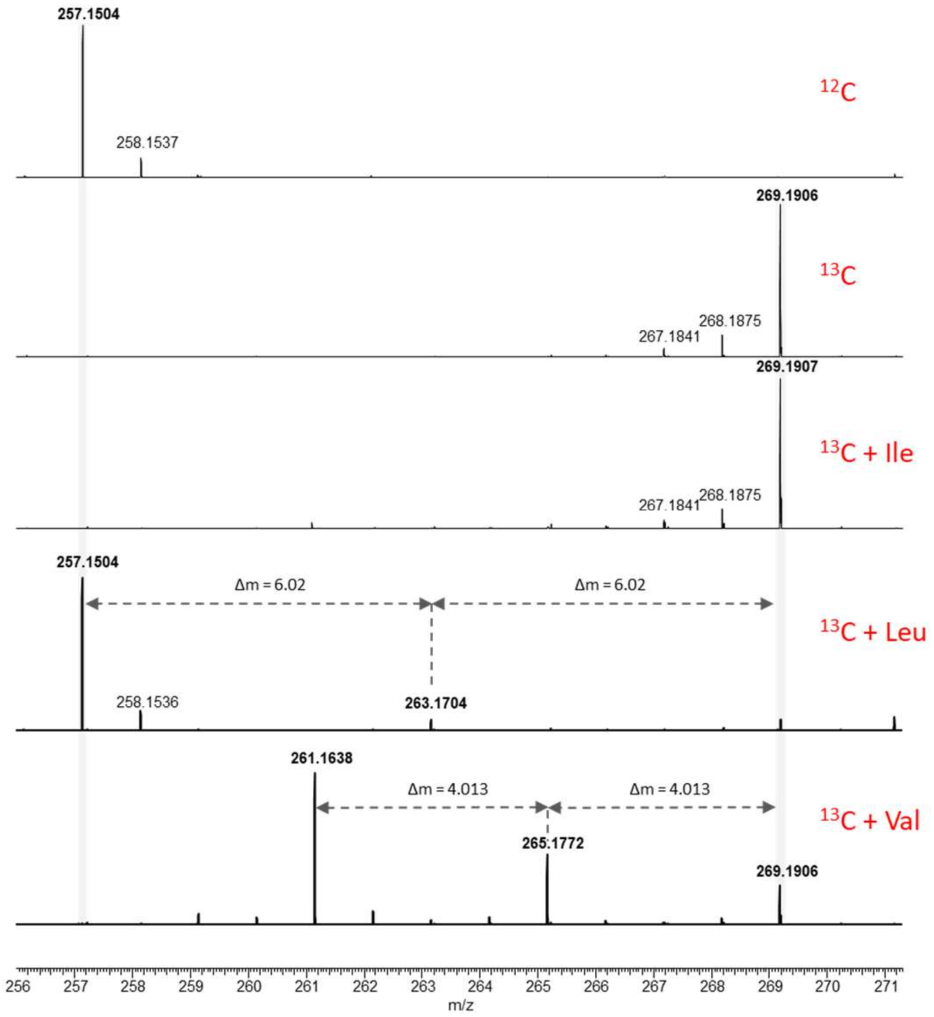


**Supplementary Figure S3:** Reverse stable isotope labeling in a $^{13}C$-background with addition of 2 mM isolecuine, leucine and valine respectively. A, B, C, E) Mass shifts show the incorporation of two leucine [2 x $C_6$] for (**1**, 227.176 *m*/*z* $[M+H]^+$, $C_{12}H_{22}N_2O_2$) confirming the diketopiperazine Cyclo(Leu-Leu), (**4**, 275.161 *m*/*z* $[M+H]^+$, $C_{12}H_{22}N_2O_5$ ), (**3**, 259.165 *m*/*z* $[M+H]^+$, $C_{12}H_{22}N_2O_4$) and (**2**, 257.150 *m*/*z* $[M+H]^+$, $C_{12}H_{20}N_2O_4$). D) Shows incorporation of a $C_6$- and $C_5$-backbone into (**5**, 229.155 *m*/*z* $[M+H]^+$, $C_{11}H_{20}N_2O_2$) both originating from leucine. For all ions, incorporation of two $C_4$-backbones is observed for valine, which is probably metabolized via 2-isopropylmalate into leucine.

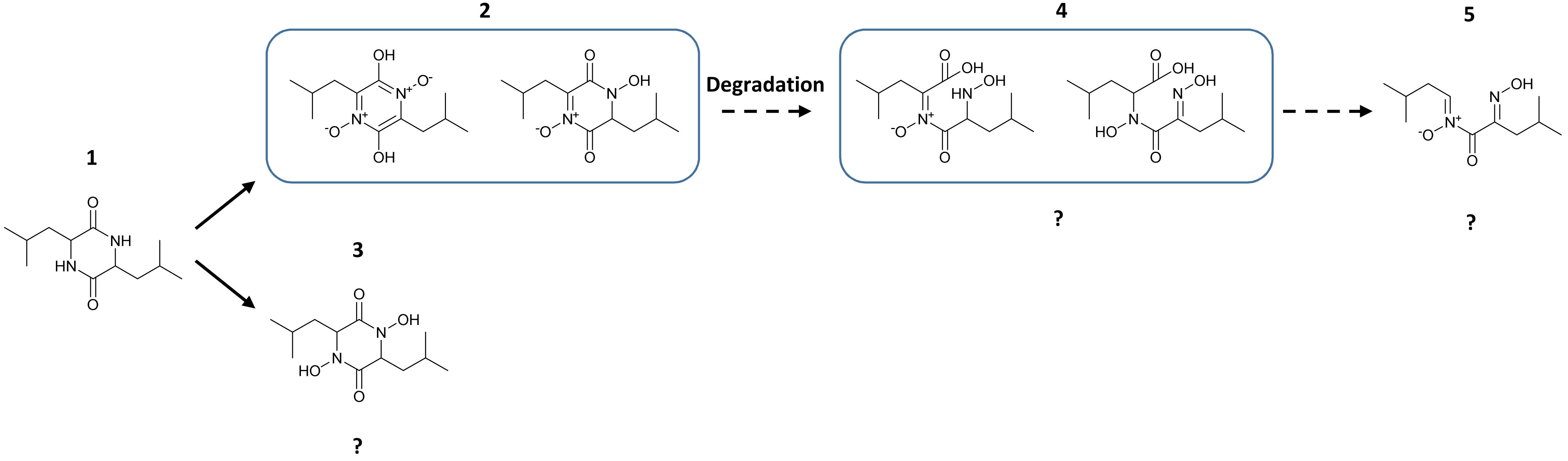


**Supplementary Figure S4:** Proposed structures for **1** Cyclo(Leu-Leu) derived metabolites. **2** pulcherriminic acid and **3** are precursors for pulcherrimin. In contrast and based on our feeding experiments **4** and **5** might represent so far undescribed degradation products of **2**.

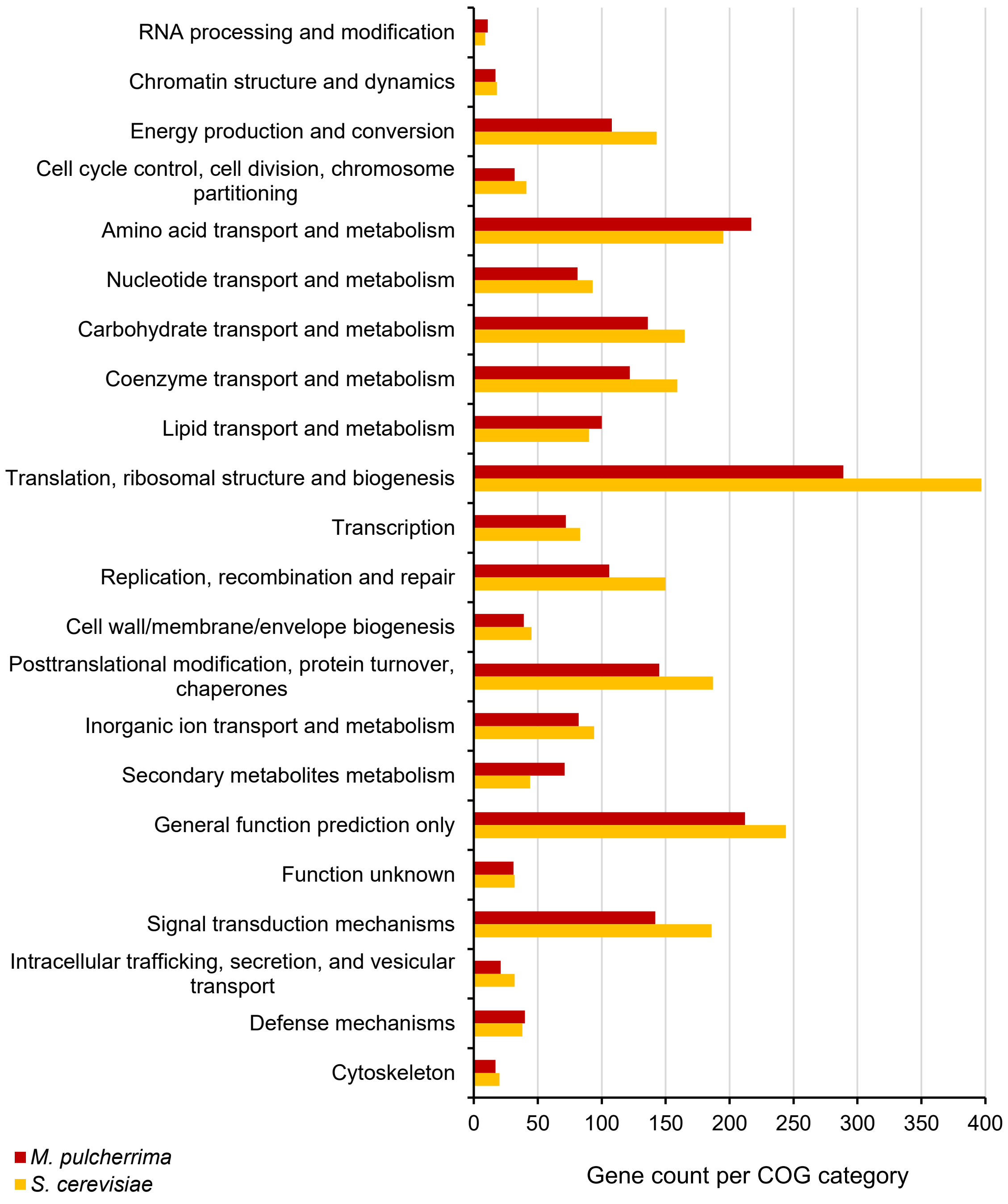


**Supplementary Figure S5**: Comparison of gene counts for different clusters of orthologous group (COG) categories in S. cerevisiae S288C (yellow) and M. pulcherrima APC 1.2 (red). The analysis was performed using the Integrated Microbial Genomes (IMG) system {Chen, 2017 #5558}.

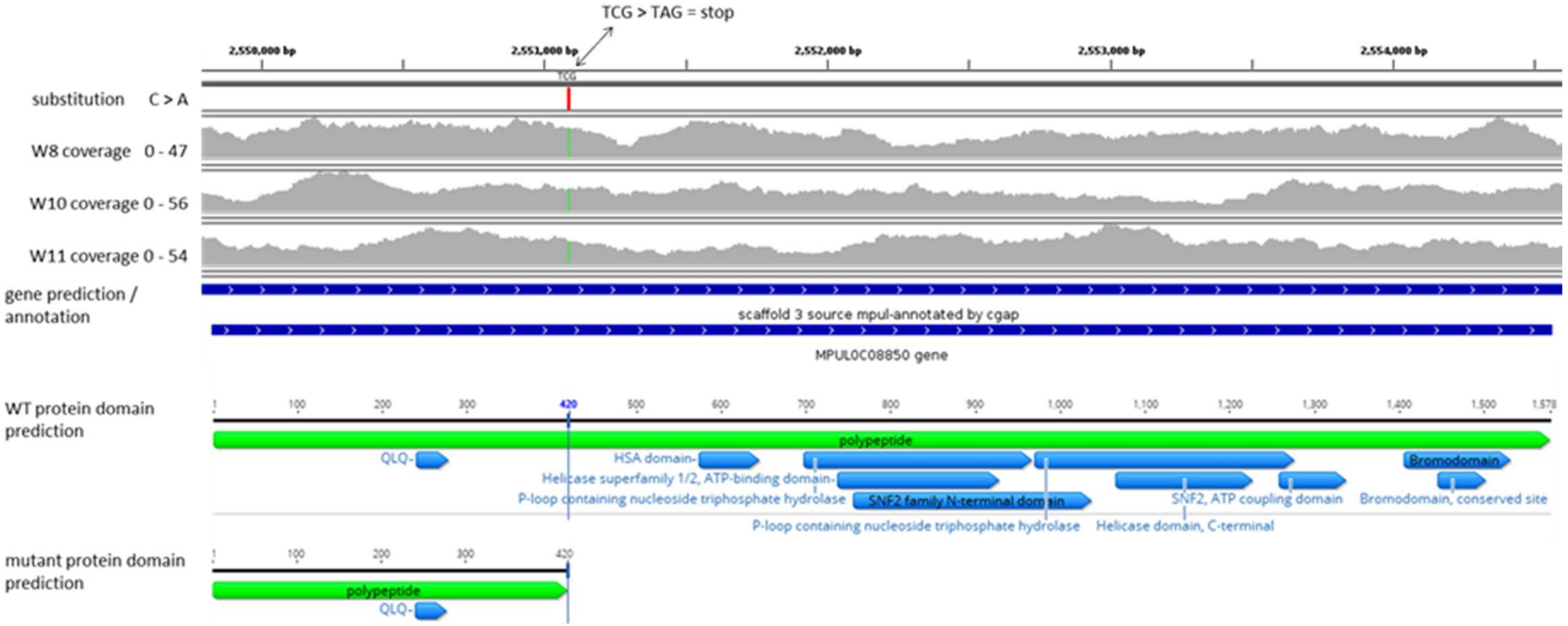


**Supplementary Figure S6: Pigmentless *M. pulcherrima* mutants harbor a point mutation leading to a premature stop codon in the *SNF2* gene.** The C → A point mutation at the nucleotide position 1262 of the Mpul *SNF2* gene (Mpul 0C08850) was predicted to result in a truncated protein of 420 amino acids in length, as compared to 1578 amino acids for the full length protein, that lacked all domains and motifs except for the first glutamine-leucine-glutamine (QLQ) domain.

*Mpul SNF2*

**A**

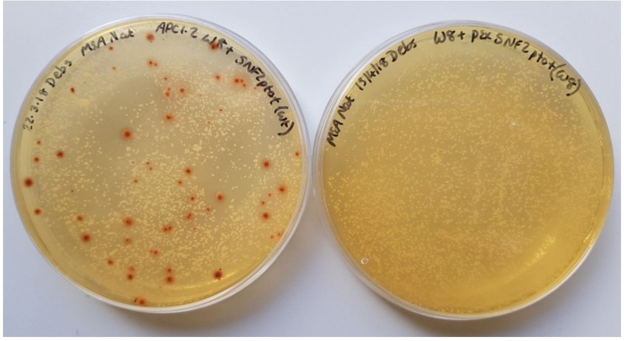

**B**

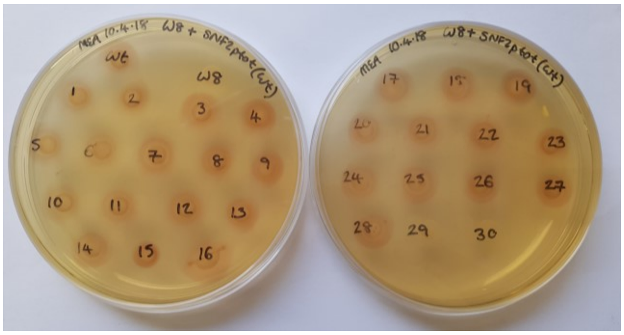


**Supplementary Figure S7**: Complementation with wildtype *SNF2* restores red pigmentation to the pigmentless *M. pulcherrima* mutant. Although *M. pulcherrima* can be transformed with high efficiency, targeted genomic integration of constructs is currently limited due to very low rates of homologous recombination, and tools for alternative approaches to genome targeting are not available at present nor are stable replicating plasmids. As such, we introduced the *SNF2* clone under the control of its native promoter into the W8 genome via random genomic integration. A) W8 mutants transformed with the wildtype *SNF2* clone (left) had red pigmentation restored, whereas those transformed with the mutant *snf2* clone (right) remained pigmentless. B) Of the thirty positive colonies picked from the *SNF2* plate, twenty-eight colonies were pigmented whilst two remained pigmentless. A possible explanation for variation in pigmentation restoration is an effect of the integration site influencing expression of the construct.
